# When Less Is Enough: Low-Rank Structure in DNA Sequence-to-Function Models

**DOI:** 10.64898/2026.01.21.700827

**Authors:** Elizabeth Gilfeather, Maria Chikina, Dennis Kostka

## Abstract

**Motivation:** The rapid success of deep learning sequence-to-function (S2F) models has driven a trend toward ever larger architectures for regulatory genomics. While these models achieve meaningful predictive performance, their growing size and computational cost have made them increasingly opaque, difficult to deploy, and impractical for widespread use. As S2F models mature, an open question is how much model capacity is truly necessary to represent regulatory sequence function.

**Results:** We show that, within linear layers, much of the predictive power of state-of-the-art S2F models is concentrated in low-rank structure. Using post hoc singular value decomposition, we construct low-rank approximations of the Sei model that reduce model size by up to 90% while preserving over 90% correlation with full-model predictions. Strikingly, extremely low-rank models—including rank 1—can match or exceed full-model performance on several variant-effect benchmarks. This indicates dominant regulatory signals may lie in a surprisingly low-dimensional subspace. Combining low-rank approximation with static model quantization enables practical CPU inference, yielding over 5× speedups relative to the full Sei model and more than 100× faster inference than larger S2F models on comparable tasks. Applying the same approach to Borzoi, Enformer, and AlphaGenome reveals consistent behavior, demonstrating that low-rank structure is a general property of modern S2F architectures, not a model-specific artifact.

**Availability and Implementation:** Low-rank Sei is available at https://github.com/kostkalab/seillra.

## 1. Introduction

Understanding how genetic variation contributes to complex traits and disease remains a central challenge in human genomics. While genome-wide association studies (GWAS) and quantitative trait locus (QTL) analyses have identified thousands of associated loci across diverse phenotypes (most in non-coding regions), the mechanisms by which non-coding variants perturb gene regulation remain difficult to resolve. Progress is limited by linkage disequilibrium, polygenicity, and the practical difficulty of experimentally assaying all relevant cell types and regulatory contexts [Visscher et al., 2017]. This motivates *in silico* approaches for predicting regulatory effects of sequence variation at scale.

Deep learning sequence-to-function (S2F) models have emerged as powerful representations of regulatory sequence, enabling the prediction of functional genomic readouts (ATAC-seq, DNase-seq, ChIP-seq, etc.) directly from DNA sequence. Models such as Sei [Chen et al., 2022], Borzoi [Linder et al., 2025], Enformer [Avsec et al., 2021], and AlphaGenome [Avsec et al., 2026] achieve strong predictive performance across assays and cell types [Benegas et al., 2025], and enable analysis of rare and *de novo* variants without requiring population-level observations. However, these models are large and computationally intensive, making their use increasingly dependent on specialized GPU hardware and limiting accessibility for exploratory analysis and large-scale variant screening. As a result, researchers often favor simpler statistical approaches despite the greater expressivity of deep learning models [Fusco et al., 2025].

One response to these challenges has been the development of intrinsically interpretable or explicitly constrained models (Balcı et al. [2023], Novakovsky et al. [2023], Tseng et al. [2024], etc.), which aim to encode regulatory logic in transparent, biologically motivated representations. While such models can provide valuable mechanistic insight, they tend to trade off predictive breadth or flexibility. In parallel, model compression techniques have been proposed to reduce the computational cost of deep learning models, including pruning [Hegde et al., 2025], knowledge distillation [Avsec et al., 2026], and low-rank adaptation via parameter-efficient finetuning [Hu et al., 2021, Makhdoomi and Ghosh, 2024]. In genomics, however, these approaches have largely been underutilized or treated solely as engineering optimizations.

Here, we take a complementary perspective by using low-rank compression of pretrained S2F models to ask how predictive performance changes as model capacity is systematically altered. Using post hoc singular value decomposition of linear layers, we construct a family of low-rank approximations for four S2F models that allows model rank to serve as a controlled axis of expressivity. In this way, we use model compression to investigate how much task-relevant signal is retained under low-rank approximation.

The resulting low-rank models are substantially smaller than their full counterparts, while maintaining high agreement with full-model predictions. When combined with static quantization [Kulkarni et al., 2022, Or et al., 2025], they enable fast, memory-efficient inference on CPUs. Across benchmarks of promoter eQTLs and cell-type-specific chromatin accessibility QTLs, the compressed models match or exceed full-model performance while dramatically reducing computational requirements. We release these models through the `seillra` Python package, providing efficient and accessible sequence-to-function models for regulatory genomics analyses.

## 2. Results

### Low-Rank Approximation of Linear Operations

We analyze four S2F models with publicly available weights: Sei [Chen et al., 2022], Borzoi [Linder et al., 2025], Enformer [Avsec et al., 2021], and AlphaGenome [Avsec et al., 2026], which predict functional sequence annotations from input DNA, but differ in architecture, sequence length, and output tracks. In all four models, a substantial fraction of parameters resides in linear layers or in modules containing linear operations, such as multi-head attention (Table 1). A standard linear layer/operation is described by:

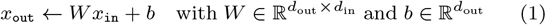

requiring *d*_out_(*d*_in_ + 1) parameters. We replace each linear layer with a rank-*k* factorization:

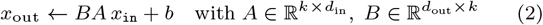

requiring only *k*(*d*_out_ + *d*_in_) + *d*_out_ parameters. When *d*_in_ and *d*_out_ are large (tens of thousands, as in our applications), even moderate *k* yield substantial parameter savings.

**Table 1.** Sequence-to-function models in this study.

| model | input length [kb] | output length [kb] | # parameters [M] | % linear | # data tracks | citation |
| --- | --- | --- | --- | --- | --- | --- |
| Sei | 4 | 0.001 | 890 | 91.7 | 21,907 | [Chen et al., 2022] |
| Enformer | 197 | 114.688 | 246 | 70.7 | 5,313 | [Avsec et al., 2021] |
| Borzoi | 524 | 196.608 | 186 | 67.8 | 7,611 | [Linder et al., 2025] |
| AlphaGenome | 4 to 1048 | 4 to 1048 | 450 | 42.6 | 5,930 | [Avsec et al., 2026] |

For the four already-trained models, singular value decomposition (SVD) [Golub and Kahan, 1965] provides a principled way to derive *A* and *B* from *W*. The optimization problem

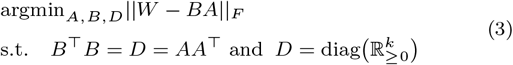

for diagonal *D* is solved by

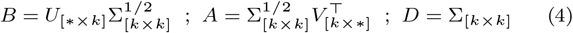

where *U* Σ*V* ^*⊤*^ is the SVD of *W* and subscripts denote appropriate sub-matrices. This approach provides a straightforward means for parameter reduction in already-trained models.

Using this approach, we replaced eligible linear operations, including those within attention blocks, with rank-k approximations for k ranging from 1 to 2,048. An operation was replaced only when its low-rank factorization required fewer parameters than the original operation. We refer to these as Sei-LLRA, Borzoi-LLRA, Enformer-LLRA, and AlphaGenome-LLRA. At rank 1, model size reductions were 91.6% for Sei, 70.6% for Enformer, 67.7% for Borzoi, and 42.5% for AlphaGenome (Fig. 1A). Multiply-accumulate operations (MACs) decrease quickly with rank; Borzoi, Enformer, and AlphaGenome require greater than 10*×* more MACs per input sequence than Sei due to their longer input and output sequences (Fig. 1B). Across architectures, the correlation between LLRA and full-model outputs increased with rank for ENCODE cCRE track predictions [The ENCODE Project Consortium, 2012], with ranks of 128 and above maintaining *>*90% correlation while reducing model size by more than 50% (Fig. 1C). Across models and ranks, low-rank predictions of DNase-seq tracks tended to show higher correlations with full-model outputs than predictions of RNA-seq tracks (Fig. 1D,E).

**Fig. 1:**
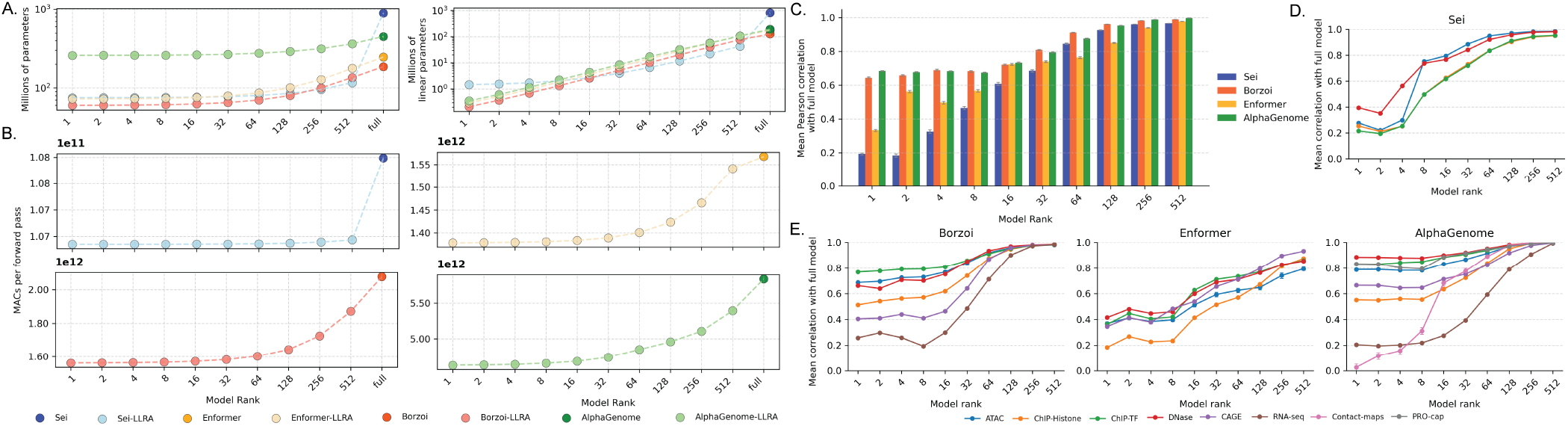
Low-rank Sei, Borzoi, Enformer, and AlphaGenome architectures efficiently compress full model architectures. **(A)** Model rank (x-axis) compared to total number of model parameters and number of linear layer parameters. **(B)** Model rank compared to number of multiply-accumulate operations (MACs). **(C)** Mean Pearson correlation between full-model and low-rank outputs across sequences centered on cCREs. **(D**,**E)** Mean Pearson correlation between full-model and low-rank outputs seperated by output modality.

### Low-rank models predict promoter variant effects

We evaluated the LLRA models on zero-shot prediction of promoter variant effects using seven benchmark variant sets within *±*500bp of protein-coding TSSs (PromoterAI benchmarks, Jaganathan et al. [2025]).

The benchmarks are either reporter assays that probe local promoter grammar (CAGI5 Saturation, MPRA Saturation, MPRA eQTLs) or natural in vivo datasets reflecting variants in their endogenous genomic context (GTEx Outliers, GTEx eQTLs, UKBB Proteome, GEL RNA). Together, these datasets provide a structured test of promoter regulation ranging from local isolated effects to context-dependent variation.

For Sei-LLRA, models with rank 64 and higher perform comparably to the full model across all datasets (Fig. 2B). Unexpectedly, the rank 1 model achieves higher auROC than the full Sei model on most datasets, whereas intermediate ranks (2–32) show reduced performance for several comparisons. This non-monotonic relationship indicates that a very low-dimensional representation is sufficient to capture much of the signal relevant for generic promoter variant effects.

**Fig. 2:**
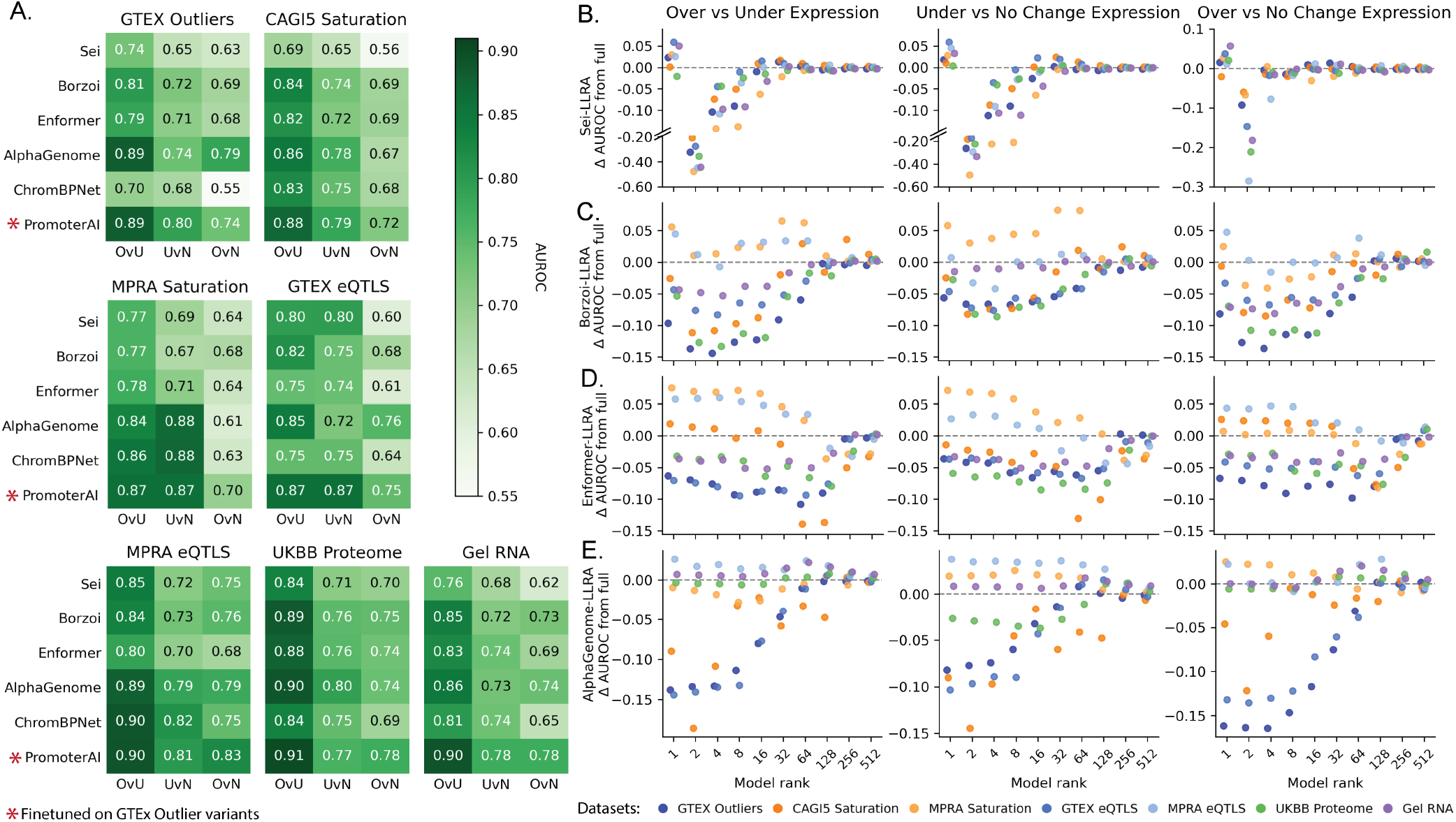
Zero-shot prediction of promoter eQTL effect-sizes with full models and low-rank approximations. **(A)** AuROC for each dataset from Jaganathan et al. [2025] for the full Sei, Borzoi, Enformer, and AlphaGenome models, and ChromBPNet and PromoterAI. Variants are classified as having the effect of over-expression, under-expression or no change. Note that PromoterAI is finetuned on a subset of variants from the GTEx Outlier dataset. **(B–E)** For Sei, Borzoi, Enformer, and AlphaGenome, the difference between each LLRA model’s and its respective full model’s prediction of the effect direction for QTL variants is shown. The y-axis shows the difference in auROC between the full model and the low-rank model.

Borzoi-LLRA and Enformer-LLRA show similar patterns (Fig. 2C,D). For Borzoi, rank 128 and above matches full model performance, with rank 1 achieving higher auROC than the full model on two datasets. For Enformer, the rank associated with the highest performance depended on the task: low ranks perform best on saturation mutagenesis datasets, while higher ranks are needed for QTL prediction. This task-dependence likely reflects differences in the sequence features relevant to each benchmark.

For AlphaGenome-LLRA, promoter variant datasets were scored using the modality that had the highest performance for the full model. Datasets scored using RNA-seq (GTEx and CAGI5) have reduced performance at low ranks, and they recover to the full model predictions by rank 128 (Fig. 2E). However for the remaining datasets scored using DNase-seq and CAGE-seq, low rank models (including rank 1) approximate or surpass full model predictions.

Across all seven datasets, LLRA versions of each model matched or exceeded performance of full models, ChromBPNet [Pampari et al., 2025], and PromoterAI (which was finetuned on GTEx eQTL outliers) [Jaganathan et al., 2025] for at least one rank (Fig. 2A–E). Across datasets and models, rank 128 appears to be a good trade-off between consistent predictive accuracy and model size reduction. These results demonstrate that low-rank approximation preserves predictive accuracy while substantially reducing model size.

### Increased inference speed and accessibility through static model quantization

Low-rank approximation reduces memory requirements, but inference on CPU hardware remains slow. Static quantization reduces memory use and may accelerate supported operations by representing model weights and activations in lower-precision integer formats, reducing memory footprint and enabling faster computation. We applied static quantization to the LLRA models, and find that model quantization substantially accelerates Sei-LLRA inference on CPUs.

Quantization reduced Sei-LLRA inference time from 1.33 to 0.34 sec/sequence, but is not meaningfully reduced for quantized Borzoi-LLRA, Enformer-LLRA, or AlphaGenome-LLRA, which remain significantly slower than the quantized Sei-LLRA model (Fig. 3A). When inference times are normalized by input length, quantized Sei-LLRA and Borzoi-LLRA both have similar speeds at less than 0.1 seconds per input-kb (Fig. 3B).

**Fig. 3:**
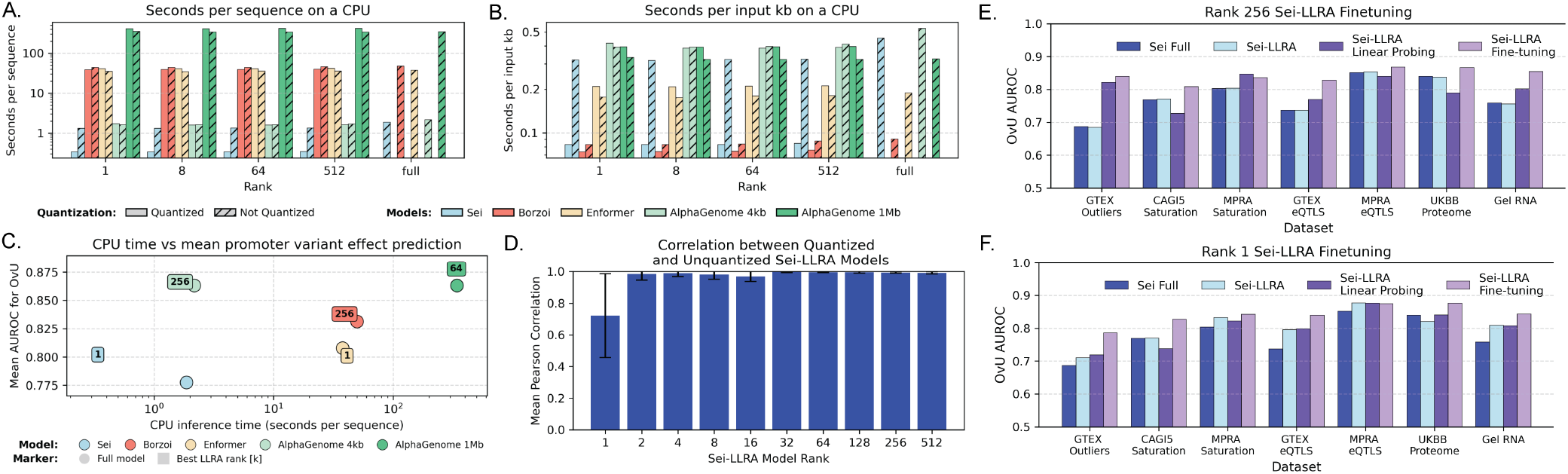
Model quantization increases inference speed and enables model finetuning of low-rank Sei on the CPU. **(A, B)** Quantized and un-quantized LLRA models and un-quantized full-rank models’ inference speed on a single thread CPU, measured in seconds per input-sequence and in seconds per input-sequence-kb. **(C)** LLRA models’ CPU inference times per sequence (x-axis) compared to mean auROC of the over vs under expression promoter variant benchmark datasets (y-axis). The rank with the highest mean auROC for each model is annotated. **(D)** Correlation between quantized and un-quantized Sei-LLRA models’ outputs for sequences centered on ENCODE cCRE elements. **(E, F)** AuROC for the prediction of over-vs. under-expression of promoter eQTL datasets. Comparison between full Sei and Sei-LLRA (ranks 1 and 256), and between linear probing and finetuning of Sei-LLRA.

However, for use cases requiring predictions across many sequences, absolute inference time matters most, and only quantized Sei-LLRA models appear practical for CPU deployment (Fig. 3A,C). We observe that quantized Sei-LLRA models’ outputs correlate well with outputs from un-quantized models on ENCODE cCRE sequences [The ENCODE Project Consortium, 2012] (Fig. 3D).

### Finetuning of Sei-LLRA on CPUs

To demonstrate that Sei-LLRA models can be adapted to specific tasks, we performed linear probing and finetuning on CPUs using outlier promoter eQTLs from Jaganathan et al. [2025]. For linear probing, Sei-LLRA sequence embeddings were used to train a small two-layer network to predict variant effect z-scores. For fine-tuning, Sei-LLRA convolution layers (quantized, frozen) produced embeddings that were passed through the model head and sequence class projection layer(un-quantized, frozen) which were then finetuned. Both procedures completed in approximately one hour using 20 CPU cores. For the rank 1 and rank 256 models, finetuning improved auROC across benchmarks, while linear probing showed limited improvement (Fig. 3E), demonstrating that Sei-LLRA models can be efficiently finetuned on CPUs to achieve meaningful gains in predictive accuracy.

### The seillra package

To make Sei-LLRA models broadly accessible, we developed two Python packages. The seimodel package (https://github.com/kostkalab/seimodel) provides modular access to the original Sei model, separating it into a sequence embedding trunk, an assay prediction head, and a sequence class projection layer. The `seillra` package (https://github.com/kostkalab/seillra) provides analogous access to quantized and non-quantized Sei-LLRA models across a selection of ranks, for CPU or GPU inference. Both packages include installation instructions and usage examples.

### Evaluation of Sei-LLRA on cell-type specific QTLs

To assess whether Sei-LLRA models capture cell-type specific regulatory variation, we evaluated zero-shot predictions on context specific QTL benchmarks, including caQTLs and dsQTLs from three human populations measured in GM12878 cells, caQTLs in specific to microglia or smooth muscle cells, and SPI1 transcription factor binding QTLs (bQTLs) in GM12878 cells [Pampari et al., 2025].

For GM12878 caQTLs and dsQTLs, Sei-LLRA models achieved similar average precision for predicting unsigned effect sizes and Pearson correlation for signed effect sizes (Fig. 4A) to the full Sei model. For caQTLs in microglia and smooth muscle cells, ranks 64 and above showed similar performance to the full model; for SPI1 bQTLs, ranks 16 and above performed similarly to full Sei (Fig. 4A,B).

**Fig. 4:**
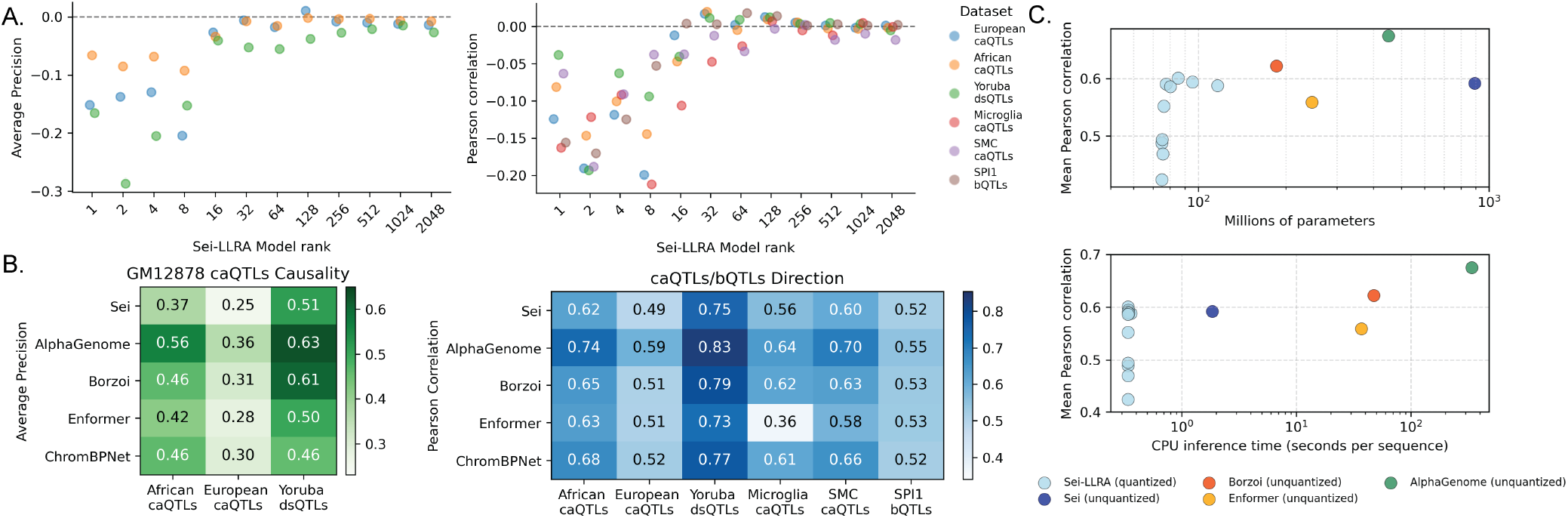
Quantized low-rank Sei models efficiently predict cell-type specific QTL effect sizes. **(A)** Difference in average precision and Pearson correlation between full Sei and low rank Sei-LLRA model predictions across causality and direction caQTL datasets from [Pampari et al., 2025]. **(B)** Full model performance across caQTL datasets for causality and direction. **(C)** Mean Pearson correlation over datasets for the seven quantized Sei-LLRA models (different ranks), and un-quantized Sei, Borzoi, Enformer, and AlphaGenome compared to each model’s number of parameters and CPU inference time.

For these data, the rank 64 Sei-LLRA model represents a good trade-off between size and performance: it typically matches full Sei accuracy while being substantially smaller. Compared to Borzoi, rank 64 Sei-LLRA shows a modest decrease in predictive performance (0.04 average correlation reduction across direction QTL datasets), but offers *>*100-fold faster CPU inference (Fig. 4C).

## 3. Discussion

We have developed compressed versions of the Sei S2F model that maintain predictive accuracy while substantially reducing computational requirements. By combining low-rank approximation of linear layers with static quantization, we reduced model size by up to 90% and achieved inference speeds of approximately .34 seconds per sequence on CPUs. Compressed Sei-LLRA models perform reasonably compared to state-of-the-art methods across variant effect benchmarks, making accurate sequence predictions accessible to researchers without large GPUs. For a starting point in practice, we recommend the use of rank 128, which offers substantial compression and speedup while maintaining good performance in our experiments. However, we expect the most reasonable choices for model rank will be data/experiment-dependent in general.

Notably, rank 1 models achieved higher auROC than full models on several promoter eQTL datasets across architectures.

This suggests that different ranks capture different types of sequence information, with varying relevance depending on the prediction task. SVD decomposes weight matrices by variance explained rather than by task relevance, and so information captured at intermediate ranks may be detrimental to prediction in isolation, but contributes positively when combined with higher-rank components. Investigating what features are captured at different ranks could yield insights into model behavior, but is beyond the scope of this work.

For Sei but not other architectures, quantization preserved prediction accuracy while substantially accelerating inference for practical CPU usage. This is most likely primarily due to the other architectures’ use of interleaved linear layers that require repeated quantization and de-quantization throughout operations like `softmax` and `gelu`, which offset efficiency gains.

We evaluated these models in the context of variant effect prediction, but low-rank approximation and quantization apply equally to other sequence-to-function tasks such as sequence embedding, motif analysis, or regulatory element classification.

In summary, the `seillra` package provides efficient, accessible Sei models for chromatin accessibility and variant effect prediction. These models enable researchers to perform sequence-to-function analyses on standard hardware, lowering barriers to applying sequence-to-function models in regulatory genomics.

## 4. Methods

### Model Compression using Singular Value Decomposition

To create low-rank version of the Sei [Chen et al., 2022], Borzoi [Linder et al., 2025], Enformer [Avsec et al., 2021], and AlphaGenome [Avsec et al., 2026] models, we replaced each of the linear layers (and layers that contain linear operations, like multi-head attention) with a low-rank approximation layer, keeping bias terms where applicable.

We downloaded the Sei model architecture from https://github.com/Function-Lab/sei-framework, and the model weights and sequence class projection matrix from https://zenodo.org/records/4906997. We downloaded the gReLU [Lal et al., 2024] implementation of the Borzoi and Enformer model architectures from https://github.com/Genentech/gReLU/tree/main/src/grelu (human “replicate 0” for Borzoi and “human” for Enformer). AlphaGenome weights were downloaded from https://github.com/genomicsxai/alphagenome-pytorch. For each model and each rank (*k* = 2^*n*^), we replaced all linear layers with larger dimensions than *k* than with low-rank layers. We calculated the Pearson correlation between the full and low-rank models prediction output for 250 sequences centered on putative cis-regulatory elements annotated by ENCODE [The ENCODE Project Consortium, 2012] (Fig. 1C-E), and report the average across sequences.

### Model Quantization

We used static quantization on the full and low rank models using `torch.ao` default quantization with the `fbgemm` mapping. For Sei, we separated the model into its 2 blocks, one with convolution layers and one with linear layers, and quantize each separately. The Sei blocks were all calibrated using 5, 000 random one-hot encoded sequences, and evaluated on the same set of random sequences. We note that quantized versions of Borzoi, Enformer, and AlphaGenome (4kb and 1Mb input, only 128bp res output) were only used for assessment of inference speed and not prediction evaluation.

We computed the inference time for each model by timing inference for 15 random sequences passed individually to the model. A mean inference time was taken across replicates. CPU inference was evaluated using a single thread and single core, on an Intel Xeon Gold 6226R (2.9Ghz, SMT disabled) CPU.

### Promoter QTL variants

Variants and labels (over-expression, under-expression, no change) for 7 QTL and MPRA datasets were downloaded from https://github.com/Illumina/PromoterAI/tree/master/data/benchmark [Jaganathan et al., 2025]. For each variant, reference and alternative allele sequences centered on the variant were generated using the hg38 genome. For Sei and the Sei-LLRA models, each sequence and its reverse complement were passed through the model and averaged, with variants scored as ALT-REF for the “Promoter” sequence class. For Enformer and Borzoi, only the forward sequence was used, following the scoring procedure of Jaganathan et al. [2025]. For AlphaGenome, only the forward sequence was used, and three scoring methods were computed: mean over the center 512bp across DNase tracks, mean over the center 512bp across CAGE tracks, and mean over the nearest gene’s exons across RNA-seq tracks. Local-context datasets (CAGI5 Saturation, MPRA Saturation, MPRA eQTLs) used 4kb inputs, while the remaining, longer-context datasets used 1Mb sequences. The reported AlphaGenome scoring method for each dataset was chosen based on the best-performing metric for the full model (RNA-seq for GTEx and CAGI5, DNase-seq for MPRA, CAGE-seq for UKBB Proteome and GEL RNA). We computed auROC for each dataset across three label comparisons: over-vs. under-expressed, under-expressed vs. no change, and over-vs. no change. ChromBPNet and PromoterAI scores were taken directly from Jaganathan et al. [2025] (Fig. 2A).

### Cell-type specific QTL

QTLs were downloaded from Pampari et al. [2025] (https://www.synapse.org/Synapse:syn59449898/files/). For each variant, we generated reference and alternative allele sequences centered on the variant using the hg38 genome, lifting over hg19-annotated variants with liftOver [Genovese et al., 2024]. Variants were filtered to chromosomes 3, 6, 9, 12, 16, 18, 19, and 21 across all datasets, consistent with Avsec et al. [2026].

For the full Sei and quantized Sei-LLRA models, forward and reverse complement sequences were both passed through the model and averaged. Predictions were then averaged over replicates of the following outputs tracks: GM12878 ATAC-seq and DNase-seq for European GM12878 caQTLs; GM12878 DNase-seq for African GM12878 caQTLs and Yoruban GM12878 dsQTLs (following Pampari et al. [2025]); Macrophage ATAC-seq and DNase-seq (closest match) for Microglia caQTLs; Smooth Muscle Cell Coronary Artery ATAC-seq for SMC caQTLs; and GM12878 SPI1 ChIP-seq for SPI1 bQTLs.

For causality and direction prediction, variant scores were compared against effect sizes as reported in Pampari et al. [2025] and Avsec et al. [2026]. Average precision and Pearson correlation for ChromBPNet and Enformer were taken directly from Pampari et al. [2025]; scores not reported there (Microglia caQTLs, SMC caQTLs, SPI1 bQTLs) were computed using `grelu` following the same approach. AlphaGenome and Borzoi metrics were taken from Avsec et al. [2026].

### Sei-LLRA model finetuning on the CPU

We performed linear probing and finetuning on two quantized Sei-LLRA models (ranks 1 and 256) using outlier variants and corresponding z-scores as outlined in Jaganathan et al.. Variants in chromosomes 8, 9, 10, 21, and 22 were excluded from training. Variants in chromosomes 10, 21, and 22 were used for validation. For linear probing, we created sequence embeddings for each variant by inputting sequences centered on the reference and alternative allele to the low-rank Sei model, and used the output *∼*22k track predictions as an embedding input to a small 2 layer network of linear layers, which was trained using MSE to predict the variant’s z-score. For finetuning, we created sequence embeddings for each variant by inputting sequences centered on the reference and alternative alleles to the quantized convolution block of the low-rank Sei model. We input this into the un-quantized low-rank linear block and sequence class projection block of the Sei model. We used a loss combining the MSE between the prediction and the variant z-score and a regularization term of the MSE between the non-finetuned low-rank model and the finetuned low-rank model’s predictions for the *∼*22k tracks.

## Code availability

Code used to produce this manuscript is available at https://github.com/egilfeather/lowrank-s2f-code.

## 6. Author contributions statement

Conceptualization: DK, MC. Funding Acquisition: DK; Investigation: EG, DK, MC; Methodology: DK, MC, EG; Software: EG, DK; Supervision: DK; Writing – original draft: EG; Writing – review & editing: DK, EG, MC.

## 7 Acknowledgments

The authors would like to acknowledge support through the University of Pittsburgh School of Medicine, and through Accenture LLP through a sponsored gift.

